# Spatial profiling of pooled mRNA-LNP delivery in vivo with NanoSTAMP

**DOI:** 10.64898/2026.08.25.746710

**Authors:** Yining Zhu, Yang Miao, Ian J. Anderson, Yuexi Li, Brandon Aghnatios, Jackie No, Jingyao Ma, Di Yu, Christine Wei, Xiaoya Lu, Jialiang Wang, Josie van de Klashorst, Hai-Quan Mao, John W. Hickey

## Abstract

Existing pooled lipid nanoparticle (LNP) screens lack spatial information on formulation localization, cellular uptake, and associated multicellular tissue responses. Here we introduce NanoSTAMP, a spatially resolved, pooled in vivo screening platform for barcoded LNP libraries that uses fluorescence in situ hybridization (FISH)-based barcode readout and is compatible with spatial omics. NanoSTAMP links LNP formulation to cell-type specific uptake, cargo expression, and nearby multicellular neighborhoods within intact tissue, which enables spatially-informed design of RNA delivery, establishing tissue architecture as a dimension of LNP performance.

## Main

Messenger RNA lipid nanoparticles (mRNA LNPs) have become a clinically validated platform, enabling several FDA-approved vaccines, while mRNA-based therapeutics have been evaluated in more than 500 registered clinical trials. Their applications are now expanding beyond vaccination to protein replacement and genome editing. These increasingly demanding applications require delivery systems to be optimized not only for total expression, but also for specific tasks in tissue distribution, cellular tropism, productive intracellular delivery and biological response.^1,2^ Their performance can be tuned through lipid chemistry, component identity, molar ratio and particle properties, which collectively influence biodistribution, cellular tropism, transfection and immune responses.^3–6^ A central challenge is therefore how to evaluate large numbers of formulations not only for whether they deliver RNA, but also for where that delivery occurs, which cells internalize and productively express the cargo, and whether those cells engage the tissue environment required for the intended therapeutic response.

Existing screening approaches trade off throughput against biological context. In vitro screens enable large libraries to be evaluated but often translate poorly in vivo^4,5,7–9^, whereas pooled in vivo screens analyzed by bulk or single-cell sequencing increase throughput but require tissue dissociation and lose spatial architecture^3,10–12^. Conversely, spatial profiling preserves anatomical localization, cellular neighborhoods and local biological responses, but typically examines one therapeutic at a time. Current approaches therefore cannot connect formulation identity at scale with tissue penetration, cellular state, neighboring populations and local responses, features that can directly inform therapeutic function. For example, vaccines may require antigen expression in activated antigen-presenting cells positioned near responsive T cells, whereas protein-replacement and genome-editing therapies must reach defined tissue compartments while minimizing delivery to off-target cell populations. Here we report NanoSTAMP, a spatially resolved pooled screening platform integrating oligonucleotide-barcoded LNP libraries, FISH-based barcoding readout, and highly multiplexed protein imaging (**Fig. 1a**). NanoSTAMP simultaneously maps formulation identity, uptake, cargo expression, antigen presentation, and delivery-associated immune responses within intact tissue.

**Figure 1.**
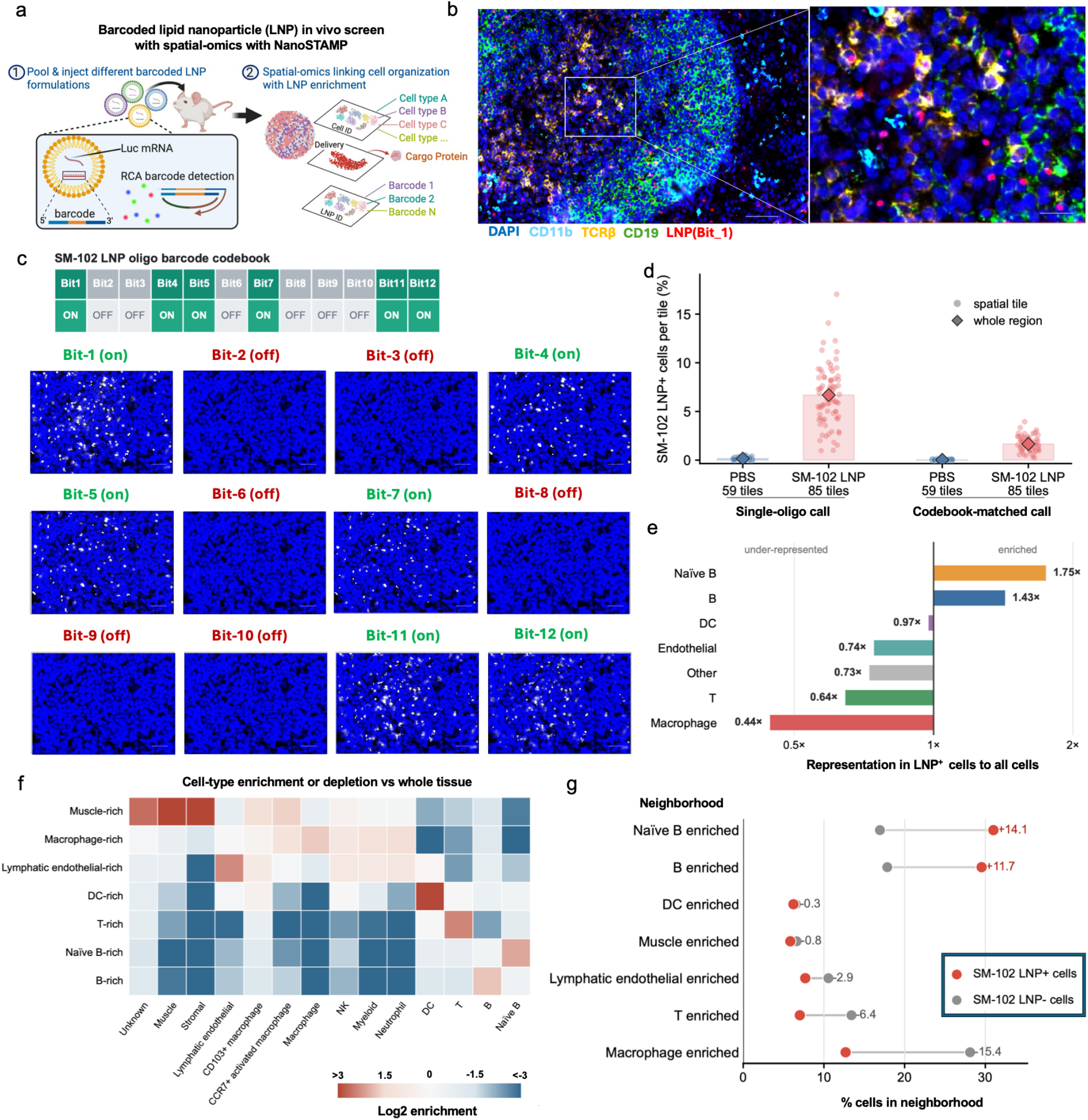
NanoSTAMP establishes spatial barcode recovery from LNP-treated spleen. **a**, NanoSTAMP Workflow for barcoded LNP delivery, RCA detection and spatial-neighborhood analysis. **b**, Representative spleen 24 h after intravenous administration of SM-102 LNPs carrying 50 μg barcode oligonucleotide and 10 μg mLuc mRNA. Scale bars, 100 μm and 25 μm. **c**, SM-102 barcode codeword and representative ON/OFF RCA channels. Scale bar, 25 μm. **d**, LNP-positive cells detected by single-bit or codebook-matched decoding across 1,500 × 1,500-pixel image tiles from the same in vivo-treated spleen section shown in b. e. **e**, Relative representation of each cell type among codebook-matched LNP+ cells compared with all cells in the same LNP-treated spleen. **f**, Spatial neighborhoods in the SM-102 LNP-treated spleen were defined by each cell’s 10 nearest neighbors and clustered by local cell-type composition using MiniBatchKMeans; heatmap shows the cell-type enrichment or depletion relative to the whole tissue. **g**, Neighborhood distributions among codebook-matched LNP-positive and LNP-negative cells. Data are representative of two experiments, each containing one PBS and one SM-102 LNP-treated spleen region.

To establish NanoSTAMP performance, B16F10 cells were treated with SM-102 LNPs used in Moderna’s COVID-19 vaccine,^1^ carrying a unique barcode oligonucleotide at 0.01, 0.1 or 1 μg ml^−1^. The barcode contained a padlock-binding site for rolling-circle amplification (RCA),^13,14^ and decoded using two fluorescent imaging probes corresponding to its two-bit codebook assignment. At 24 h, RCA recovered both probe signals at all concentrations, with minimal PBS background and greater sensitivity than direct hybridization. (**Supplementary Fig. 1**). Barcode signals also remained detectable for 72 h following treatment at 1 μg ml^−1^, supporting subsequent in vivo tracing (**Supplementary Fig. 2**). We next administered SM-102 LNPs containing 50 μg barcode oligonucleotide and 10 μg luciferase mRNA intravenously and collected spleens after 24 h. Barcode and protein signals were recovered concurrently with CODEX (CO-Detection by indEXing) multiplexed imaging,^15^ enabling LNP-positive cells to be segmented, annotated and spatially localized (**Fig. 1a, b and Supplementary Fig. 3**).

**Figure 2.**
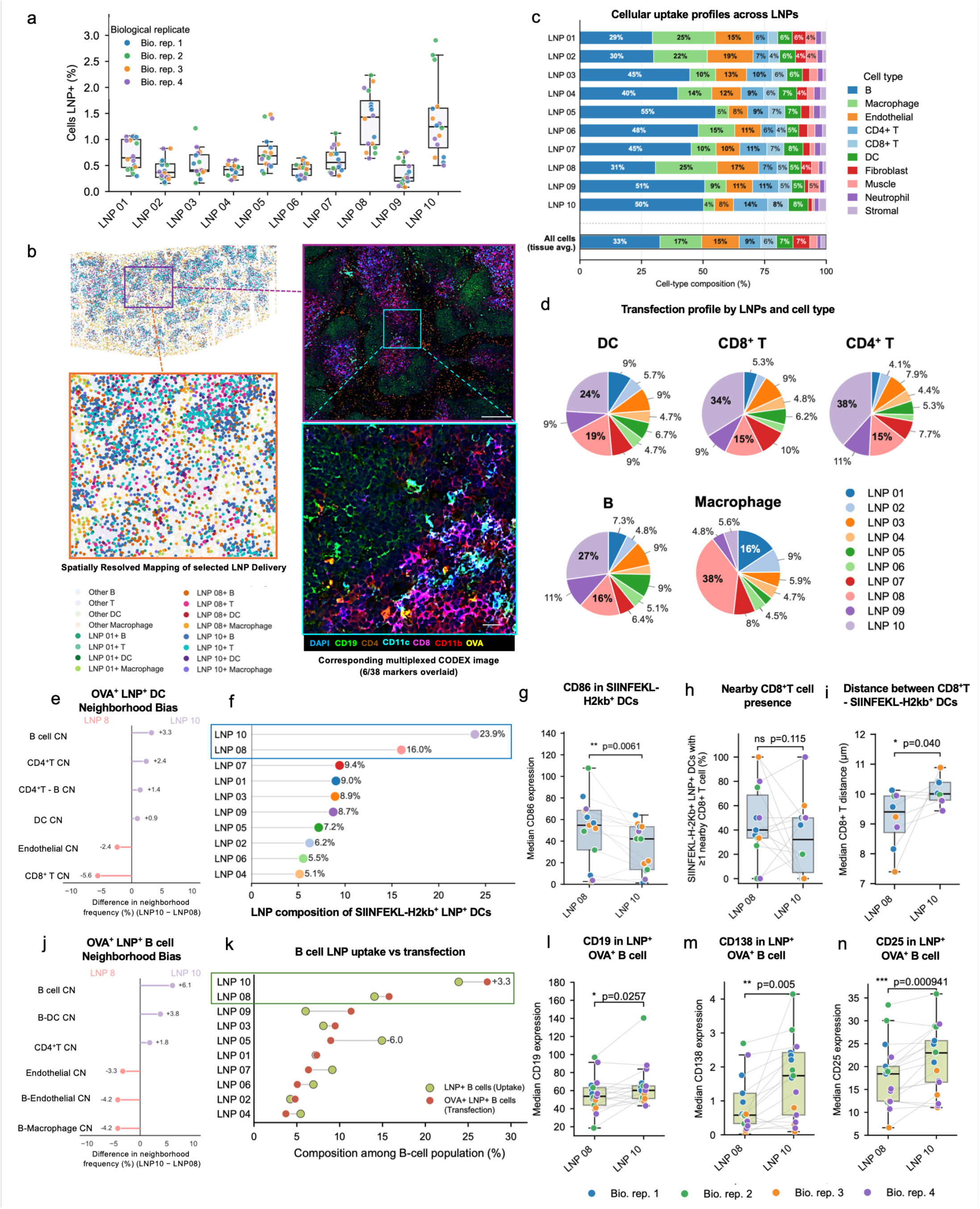
NanoSTAMP enables in vivo multiplexed profiling of LNP uptake and antigen-expression-associated cellular responses. Ten barcoded LNPs were pooled and administered intravenously at total doses of 10 μg mOVA mRNA and 20 μg barcode oligonucleotide. Spleens were collected after 24 h from four mice; four sections per mouse were independently decoded. **a**, Cellular uptake by formulation. **b**, Spatial maps of LNP01-, LNP08- and LNP10-positive B cells, T cells, DCs and macrophages, shown with CODEX imaging and OVA expression. Scale bars, 200 μm and 20 μm. **c**, Cell-type composition of LNP^+^ cells. **d**, LNP composition among OVA^+^LNP^+^ immune cells. **e**, Neighborhood bias among OVA^+^ LNP^+^ DCs based on the 20 nearest non-self cells (k=20). Values show LNP 10 minus LNP 08 percentage-point differences. **f**, LNP composition among SIINFEKL–H-2K^b+^LNP^+^ DCs. **g–i**, Paired region-level comparisons between LNP08 and LNP10: median CD86 expression across SIINFEKL–H-2K^b+^ DCs (**g**), percentage of these DCs with ≥1 neighboring CD8^+^ T cell (**h**), and median distance to the nearest CD8^+^ T cell (**i**). Each point represents a value calculated across all qualifying cells within one tissue region. **j**, Neighborhood bias among OVA^+^ LNP^+^ B cells, calculated as in i (k=20). Positive and negative values indicate LNP 10 and LNP 08 enrichment, respectively.**k**, Relationship between B-cell LNP uptake and OVA+ readout across LNP formulations. **l-n**, Paired comparisons of B-cell-associated activation and neighborhood markers between LNP 08 and LNP 10. Points represent independently processed spleen sections, colored by biological replicate. Paired lines connect matched sections. P values were calculated using two-sided paired Student’s t-tests across complete section-matched pairs. CN, cell neighborhood.

To increase multiplexing capacity, we designed a 12-bit binary codebook with Hamming weight = 6, Hamming distance ≥ 4, and applied it to the same spleen tissue section. This error-robust, codebook-matched decoding approach enables confident barcode detection in tissues while minimizing interference from spurious fluorescent signals. (**Fig. 1c, d**). Spatial neighborhoods were then defined from each cell’s ten nearest neighbors and clustered by cell-type composition. SM-102 LNP-positive cells occurred more frequently in B cell- and naïve B cell-enriched neighborhoods and less frequently in T cell- and macrophage-enriched neighborhoods (**Fig. 1e– g**). Their lower representation in macrophage-rich regions may reflect differences in tissue access, retention or phagocytic clearance.

We next applied NanoSTAMP to an in vivo screen of ten mRNA LNPs combining SM-102 or ALC-0315, the ionizable lipids used in the Moderna and Pfizer–BioNTech COVID-19 vaccines, respectively. Each was paired with a cationic (DOTAP or DDAB), zwitterionic (DSPC or DOPE) or anionic (18PG) helper lipid, selected to span lipid charge classes that we previously found to produce distinct in vivo delivery profiles.^4–6^ Each formulation carried a model ovalbumin antigen mRNA (mOVA) and a distinct 12-bit barcode (**Supplementary Fig. 4**).

The pooled library was administered intravenously, and spleens were analyzed 24 h later with CODEX multiplexed imaging. LNP08 and LNP10 reached two-to threefold more splenic cells than the DSPC-containing LNP01 and LNP06 reference formulations (**Fig. 2a, b and Supplementary Fig. 5**). The strong splenic delivery of the anionic 18PG-containing LNP10 was consistent with previous reports that anionic helper lipids can promote splenic tropism, whereas the similarly high uptake of the cationic DDAB-containing LNP08 was unexpected, indicating that helper-lipid charge alone does not fully predict in vivo delivery.^3,5^ Cellular tropism also differed: LNP05, LNP09 and LNP10 favored B cells; LNP03 and LNP07 favored T cells; and LNP01, LNP02 and LNP08 favored macrophages (**Fig. 2c and Supplementary Fig. 6**).

Productive expression was not determined by uptake alone but also by transfection efficiency and in target cell types. LNP10 was most abundant among OVA^+^LNP^+^ dendritic cells (DCs), T cells and B cells, whereas LNP08 predominated in macrophages (**Fig. 2d, Supplementary Fig. 7**). Transfection capacity, defined as the percentage of LNP^+^ cells expressing OVA, varied across formulations and cell types; LNP09 showed the strongest profile, ranging from 10–20% in lymphocytes and DCs to 35.1% in macrophages and 42.9% in non-immune cells (**Supplementary Fig. 8**). The limited correspondence between barcode recovery and OVA expression likely reflects the distinct biological steps captured by these readouts: barcode recovery reports cellular uptake, whereas OVA expression additionally requires successful endosomal escape, cytosolic mRNA release and translation. Consistent with our previous work, LNP composition and cell identity can differentially influence these processes, leading to variation in the fraction of barcode-positive cells that achieve productive expression.^4–6^

NanoSTAMP further showed that formulation effects extended beyond cellular uptake and transfection to the local neighborhoods in which these events occurred. Overall, cells with detectable LNP uptake were enriched for B-cell, neutrophil and T-cell neighbors but depleted of macrophage, fibroblast and endothelial neighbors (**Supplementary Fig. 9**). However, neighborhood distributions varied substantially across formulations: B cell-enriched neighborhoods ranged from 29.7% for LNP01 to 57.4% for LNP05, whereas macrophage-enriched neighborhoods ranged from 2.5% for LNP10 to 14.6% for LNP01 (**Supplementary Fig. 10**). The neighborhoods of cargo-expressing cells, defined by detectable LNP uptake and OVA expression, also differed across formulations and from those associated with uptake alone. B cell-enriched neighborhoods surrounding cargo-expressing cells ranged from 15.1% for LNP02 and LNP08 to 43.1% for LNP10, whereas macrophage-enriched neighborhoods ranged from 6.4% for LNP10 to 25.5% for LNP08 (**Supplementary Fig. 11**). For LNP08, B cell-enriched neighborhoods decreased from 31.4% among uptake-positive cells to 15.1% among cargo-expressing cells, while macrophage-enriched neighborhoods increased from 14.5% to 25.5%. These findings reveal an additional layer of formulation dependence: even when LNPs reached or transfected the same cell population, those cells were positioned within different cellular neighborhoods. Thus, LNP composition determines not only which cells receive and express the cargo, but also where those cells are located and which neighboring populations they may interact with, the spatial features that could shape downstream therapeutic responses.

We then focused on LNP08 and LNP10 because they were the most abundant formulations detected in dendritic cells and B cells, two cell types central to vaccine responses. A focused 20-nearest-neighbor analysis showed that OVA^+^ LNP^+^ DCs carrying LNP10 were more frequently found in B-cell- and CD4^+^ T-cell-enriched neighborhoods, whereas those carrying LNP08 were biased toward CD8^+^ T-cell-enriched neighborhoods (**Fig. 2e, Supplementary Fig. 12**). Indeed, although LNP10 represented a larger fraction of SIINFEKL–H-2K^b+^LNP^+^ DCs than LNP08 (23.9% versus 16.0%), LNP08-associated DCs expressed more CD86 (P = 0.0061), and were positioned closer to CD8^+^ T cells (P = 0.040; **Fig. 2f–i**).

Similar focused neighborhood analysis further showed relative LNP10 enrichment in B-cell-, B-cell–DC- and CD4^+^ T-cell-enriched neighborhoods, whereas LNP08 was enriched in B-cell–macrophage- and B-cell–endothelial-enriched neighborhoods (**Fig. 2j, Supplementary Fig. 13**). Moreover, LNP10-positive OVA-expressing B cells displayed significantly higher CD19, CD138 and CD25 expression than their LNP08 counterparts, consistent with a more mature and activated B-cell phenotype (**Fig. 2k–n**).

When evaluated using only the non-spatial measurements retained after tissue dissociation, cell-type-specific uptake, OVA expression and antigen presentation, LNP10 would have been prioritized for both DC- and B-cell-directed vaccination. Incorporating spatial phenotypes changed this interpretation: although LNP10 reached more antigen-presenting DCs, LNP08-associated DCs expressed more CD86 and were positioned closer to CD8^+^ T cells, suggesting that LNP08 may be better suited to support cellular immunity. In contrast, LNP10-positive OVA-expressing B cells exhibited a more differentiated and activated phenotype, supporting its potential to enhance humoral immunity. Together, these findings show how spatial context can refine LNP selection by revealing formulation-specific immune programs that are not apparent from uptake or expression alone, thereby informing the design of vaccines with different cellular or humoral objectives.

More broadly, NanoSTAMP makes delivery location and microenvironmental context actionable design variables for LNP engineering. For vaccines, it could identify formulations that deliver antigen to selected antigen-presenting cells within defined immune niches, enabling LNPs to be engineered to favor cellular or humoral immunity and potentially distinct helper T-cell programs, including Th1- or Th2-biased responses. For in vivo gene editing and protein replacement, NanoSTAMP could distinguish productive delivery to intended cell types and anatomical compartments from uptake by off-target populations, quantify editing efficiency or functional protein expression, and map local immune responses associated with toxicity or the early immune-mediated elimination of edited cells. During formulation development and manufacturing scale-up, the same approach could determine whether changes in lipid composition, particle properties or process conditions preserve the intended spatial delivery profile rather than bulk expression alone. Across these applications, the resulting multidimensional datasets could establish generalizable design principles and train machine-learning models to engineer LNPs that reach a defined cell population within a defined tissue context and elicit a desired biological function.

## Methods

### Ethical statement

All animal experiments were conducted in compliance with relevant ethical regulations and approved by the Duke University Institutional Animal Care and Use Committee under protocol A149-24-12. Male and female C57BL/6 mice aged 6–8 weeks were obtained from The Jackson Laboratory and housed under standard conditions in accordance with institutional guidelines.

### Materials

SM-102, ALC0315 was purchased from BroadPharm. DOPE, DSPC, 18PG, and DMG-PEG-2000 were obtained from Avanti Polar Lipids. Cholesterol was from Sigma-Aldrich. B16F10 (CRL-6475) were purchased from ATCC (American Type Culture Collection, USA). All mRNA (OVA mRNA and Luc mRNA) constructs were purchased from Hongene Biotech Corporation. The delivered barcode oligonucleotides and padlock probes were synthesized by Integrated DNA Technologies (IDT), and their sequences are provided in **Supplementary Table 1**. CODEX antibody information was compiled in **Supplementary Table 4**. All other chemical reagents were purchased from Sigma Aldrich (St. Louis, MO, USA) unless otherwise noted.

### LNP synthesis and characterization

LNPs were prepared by directly adding the ethanol phase to the aqueous phase at a 3:1 aqueous-to-ethanol volume ratio in 1.5-ml microcentrifuge tubes. The ethanol phase contained SM-102 or ALC-0315, cholesterol, DMG-PEG2000 and one helper lipid selected from DOTAP, DDAB, DSPC, DOPE or 18PG. Luciferase-encoding or mOVA mRNA and the corresponding barcode oligonucleotide were dissolved in 25 mM magnesium acetate buffer (pH 4.0). LNPs intended for in vivo administration were dialyzed against deionized water using a 100-kDa molecular-weight-cutoff cassette at 4 °C for 24 h and stored at 4 °C until use.

### Animals

All animal procedures were approved by the Duke University Institutional Animal Care and Use Committee under protocol A149-24-12 and performed in accordance with institutional guidelines. Male and female C57BL/6 mice aged 6–8 weeks were obtained from The Jackson Laboratory. Mice were housed in standard cages with corncob bedding under controlled conditions (18–26 °C, 30–70% relative humidity and a minimum of ten room-air changes per hour) with ad libitum access to standard chow and water. LNPs were administered intravenously at the indicated doses, and spleens were collected 24 h after injection.

### CODEX multiplexed protein and barcode imaging

Spleens were collected, embedded in optimal cutting temperature (OCT) compound and immediately frozen. Frozen tissues were sectioned at 7 μm using an Epredia HM525 NX cryostat and mounted on slides. Sections were stained with a validated panel of oligonucleotide-conjugated CODEX antibodies and imaged using successive cycles of fluorescent reporter annealing, imaging and stripping according to an established protocol.^15^ Antibody panels and imaging metadata are provided in **Supplementary Table 2**. After completion of protein imaging, slides were removed from the instrument, permeabilized and subjected to padlock-probe circularization and rolling-circle amplification to recover the delivered LNP barcodes. The resulting amplicons were decoded by sequential fluorescent-reporter hybridization and imaging according to the scheme provided in **Supplementary Table 3**. Barcode, padlock-probe and reporter sequences are provided in **Supplementary Table 1**. Raw images were processed using Akoya PhenoCycler-Fusion software (version 2.2.0) for stitching, drift correction, deconvolution and cycle concatenation. Markers exhibiting nonspecific staining or insufficient signal-to-noise were excluded before downstream analysis. Processed images were visualized using Fiji/ImageJ.

### CODEX single-cell segmentation

To obtain quantitative single-cell information, individual cells and extracted single-cell protein expression were segmented. Processed data were segmented using the SPACEc package, which can also be downloaded here (https://github.com/yuqiyuqitan/SPACEc/tree/master).^16^ SPACEc incorporates Mesmer and Cellpose, which are both deep learning-based segmentation methods.^17,18^ Mesmer was used for our segmentation, with DAPI (nuclei) along with CD45 (surface membrane) as reference channels.

### Cell-type analysis

Cells across all spleen sections were segmented using DAPI and membrane-marker signals. Segmented objects outside the predefined nuclear-intensity or size ranges were excluded, and protein-marker expression was Z-normalized before clustering. Lineage markers were used for unsupervised clustering, whereas selected functional and phenotypic markers were excluded to avoid influencing cell-type assignment. Cells were overclustered using the Leiden algorithm implemented in Scanpy. Clusters were annotated according to their marker-expression profiles and spatial correspondence with the original fluorescence images. Heterogeneous clusters were further divided or reclustered using K-means implemented in scikit-learn, and low-quality or unclassifiable objects were excluded from downstream analysis.

### Statistics & Reproducibility

Statistical analyses and data visualization were performed using custom Python scripts. Comparisons between LNP08 and LNP10 were performed using two-sided paired Student’s *t*-tests across complete section-matched pairs. Associations between LNP uptake and OVA expression were assessed using two-sided Spearman rank correlation. Statistical tests, sample sizes and exact *P* values are provided in the corresponding figure legends. *P* < 0.05 was considered statistically significant.

No statistical method was used to predetermine sample size. The multiplexed in vivo screen included four biologically independent mice, with four nonadjacent spleen sections from each mouse processed independently as technical replicates. No animals were excluded from the analysis. Low-quality or unclassifiable cellular objects were removed according to the predefined image-processing and annotation criteria described above. Blinding was not performed during sample processing because all animals received the same pooled LNP library; formulation identities were assigned computationally after imaging using the predefined barcode codebook. The single-formulation feasibility study was performed in two independent experiments with consistent results.

Fig. 1a in this manuscript were created using BioRender.com, (Created in BioRender. Zhu, Y. (2026) https://BioRender.com/z1p7jhs).

## Data Availability

### Data and materials availability

Processed data tables, cell annotations, and intermediate processed files required to reproduce the downstream figure analyses are available in the accompanying https://research.repository.duke.edu/record/554?ln=en. Raw and registered microscopy images are not included. Cell segmentation, clustering, and cell-type annotations generated using the previously published spatial-omics workflow are provided as frozen processed outputs.

### Code availability statement

Jupyter notebooks and environment requirements supporting Supplementary Figure 1c, Figure 1d–g, Figure 2, and Supplementary Figures 6–11 are available in the https://github.com/HickeyLab/NanoSTAMP. The notebooks document image stitching, spot detection, barcode matching, cell-level quantification, and downstream spatial-neighborhood analyses. Image-level processing steps require the original, non-deposited microscopy images, whereas downstream analyses can be reproduced using the deposited processed data. CODEX image segmentation was performed using the publicly available SPACEc package, which is accessible at https://github.com/yuqiyuqitan/SPACEc. SPACEc incorporates the open-source Mesmer and Cellpose segmentation algorithms.

## Author Contributions Statement

### Author contributions

Y.Z. and J.W.H. conceived of and designed this study. J.W.H. and H.-Q.M. secured the funding for this study. Y.Z., Y.M., I.J.A., Y.L., B.A., J.N., J.M., D.Y., C.W., X.L., J.W., and J.v.d.K. performed the experiments. Y.Z., Y.M., I.J.A., Y.L., B.A., and J.W.H. participated in data analysis and interpretation. The manuscript was written by Y.Z. and with revisions by J.W.H., Y.M., and inputs from all the other authors.

