## Supplemental Information for "Spatial profiling of pooled mRNA-LNP delivery in vivo with NanoSTAMP"

**Supplementary Figures**

**Figure 1.**	In vitro evaluation of NanoSTAMP detection of LNP-delivered barcode oligonucleotides.

**Figure 2.**	Time course of in vitro NanoSTAMP barcode detection following LNP delivery.

**Figure 3.**	NanoSTAMP-based spatial and phenotypic characterization of SM-102 LNP uptake in the spleen.

**Figure 4.**	Multiplexed LNP formulation and barcode codebook.

**Figure 5.**	Cell-type annotation in the multiplexed in vivo NanoSTAMP experiment.

**Figure 6.**	Cell-type-specific LNP uptake across formulations.

**Figure 7.**	Percentage of LNP-positive cells among OVA-expressing immune cells.

**Figure 8.**	Cell-type-specific LNP transfection capacity across formulations.

**Figure 9.**	Spatial cellular neighborhoods associated with decoded LNP-positive cells.

**Figure 10.**	Spatial neighborhood analysis of cells with LNP uptake.

**Figure 11.**	Spatial neighborhoods associated with LNP-mediated transfection.

**Figure 12.**	Focused spatial neighborhoods of OVA⁺ LNP⁺ dendritic cells.

**Figure 13.**	Focused spatial neighborhoods of OVA⁺ LNP⁺ B cells.

**Supplementary Table**

**Table 1.**	Delivery oligonucleotide and padlock probe sequences used in NanoSTAMP.

**Table 2.**	CODEX protein panel staining conditions and cycle information.

**Table 3.**	CODEX post-RCA staining conditions and cycle information.

**Table 4.**	CODEX antibody information.

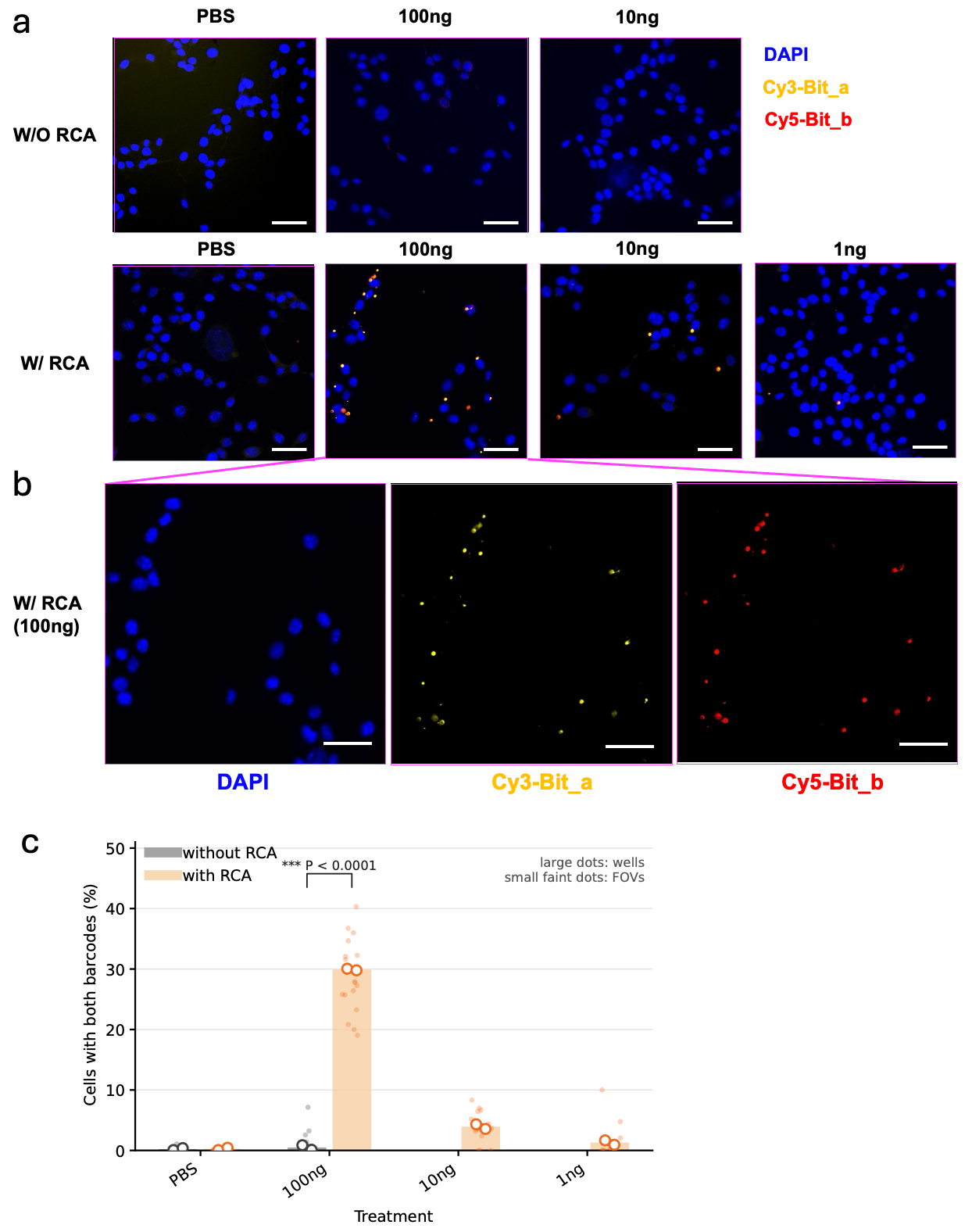

Supplementary Figure 1. In vitro evaluation of NanoSTAMP detection of LNP-delivered barcode oligonucleotides. B16F10 cells were seeded at 20,000 cells per well in 96-well plates, allowed to attach overnight, and treated with SM-102 LNPs containing a single barcode oligonucleotide sequence carrying two detectable bit regions, Bit_a and Bit_b. Doses correspond to 100, 10, or 1 ng of encapsulated barcode oligonucleotide; PBS served as a negative control. Barcode signals were assessed 24 h after treatment using direct or RCA-assisted detection. (a), Representative merged images showing nuclei (DAPI, blue), Bit_a (Cy3, yellow), and Bit_b (Cy5, red). RCA-assisted detection enabled NanoSTAMP analysis across the tested dose range, including 1 ng, with minimal background in PBS controls. (b), Enlarged single-channel images of the 100-ng condition analyzed using RCA. c, Percentage of cells positive for both Bit_a and Bit_b. Bars show group means, large open circles represent individual wells, and small faint dots represent individual fields of view. The indicated comparison at 100 ng yielded P < 0.0001. Scale bars, 50 μm.

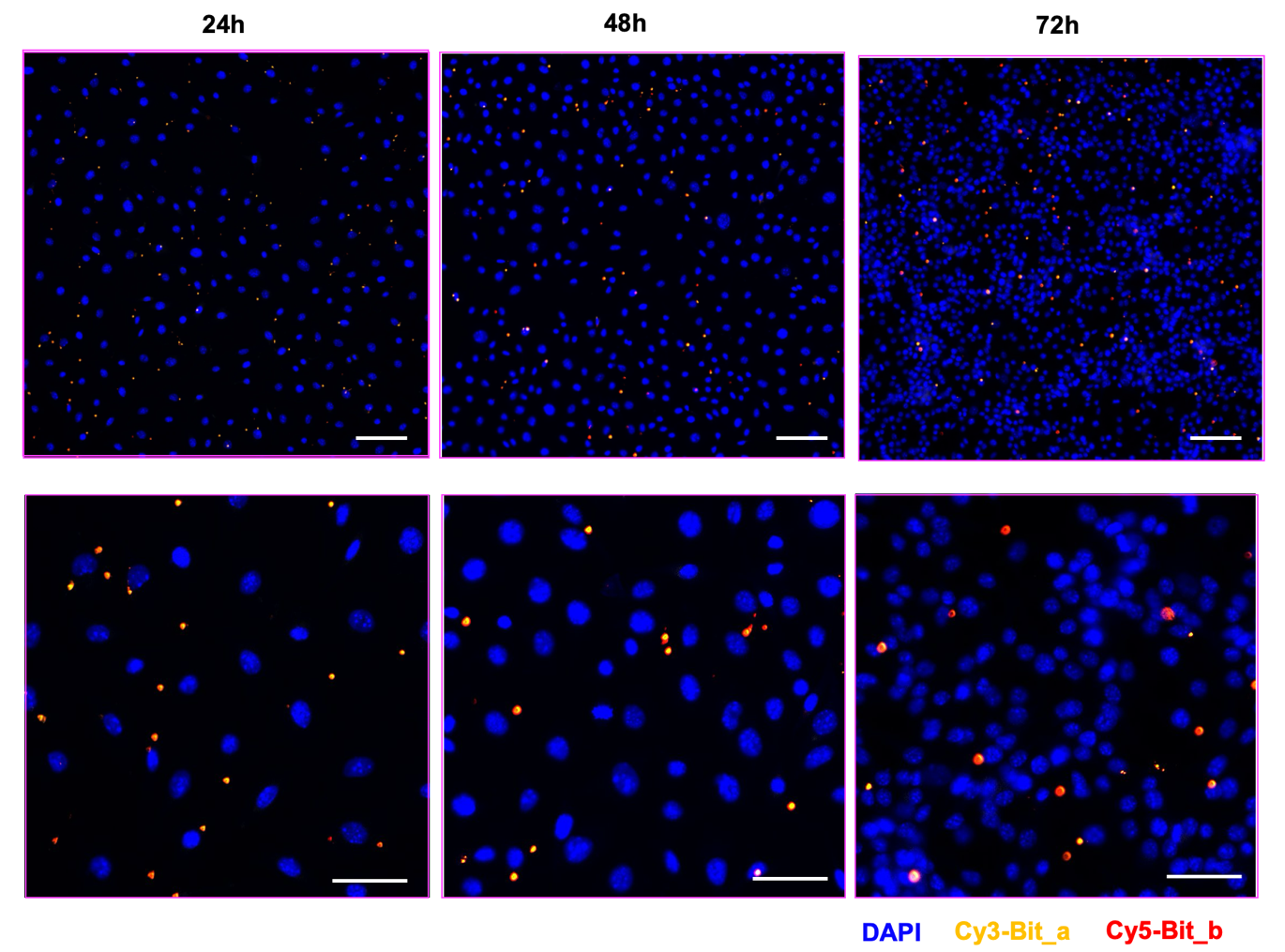

Supplementary Figure 2. Time course of in vitro NanoSTAMP barcode detection following LNP delivery. B16F10 cells were treated with SM-102 LNPs containing a single barcode oligonucleotide carrying the Bit_a and Bit_b detection regions at a dose corresponding to 100 ng encapsulated oligonucleotide. Cells were analyzed at 24, 48, and 72 h after treatment using RCA-assisted detection. Representative merged images show nuclei (DAPI, blue), Bit_a (Cy3, yellow), and Bit_b (Cy5, red). The upper and lower rows show lower- and higher-magnification views, respectively. Scale bars, 100 μm (upper row) and 50 μm (lower row).

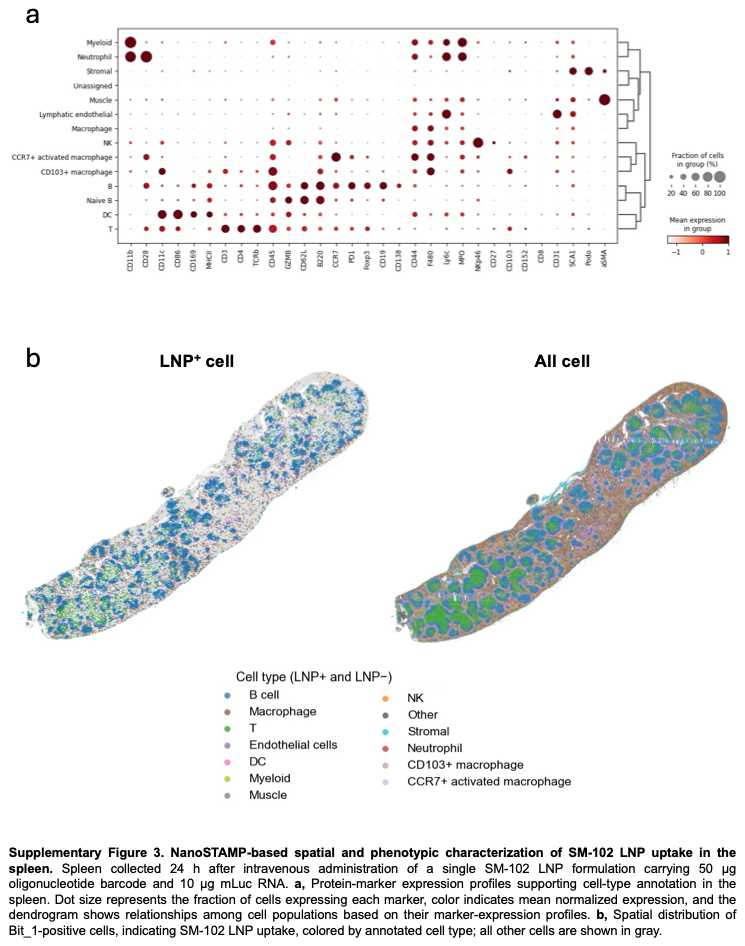

Supplementary Figure 3. NanoSTAMP-based spatial and phenotypic characterization of SM-102 LNP uptake in the spleen. Spleen collected 24 h after intravenous administration of a single SM-102 LNP formulation carrying 50 μg oligonucleotide barcode and 10 μg mLuc RNA. (a), Protein-marker expression profiles supporting cell-type annotation in the spleen. Dot size represents the fraction of cells expressing each marker, color indicates mean normalized expression, and the dendrogram shows relationships among cell populations based on their marker-expression profiles. (b), Spatial distribution of Bit_1-positive cells, indicating SM-102 LNP uptake, colored by annotated cell type; all other cells are shown in light gray.

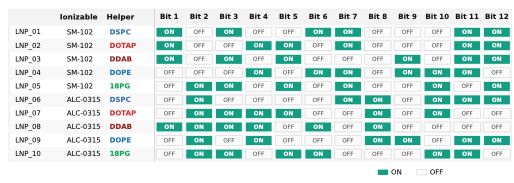

Supplementary Figure 4. Multiplexed LNP formulation and barcode codebook. Ten LNP formulations were generated using two ionizable lipids (SM-102 or ALC-0315) combined with five helper lipids (DSPC, DOTAP, DDAB, DOPE, or 18PG). Each formulation was assigned a unique 12-bit oligonucleotide barcode codeword for identification following pooled administration. Green “ON” and gray “OFF” entries indicate the presence or absence, respectively, of each barcode bit. The LNP was formulated with ionizable lipid, helper lipid, cholesterol and DMG–PEG2000 at a target molar ratio of 46.3:9.4:42.7:1.6 and an N/P ratio of 6.

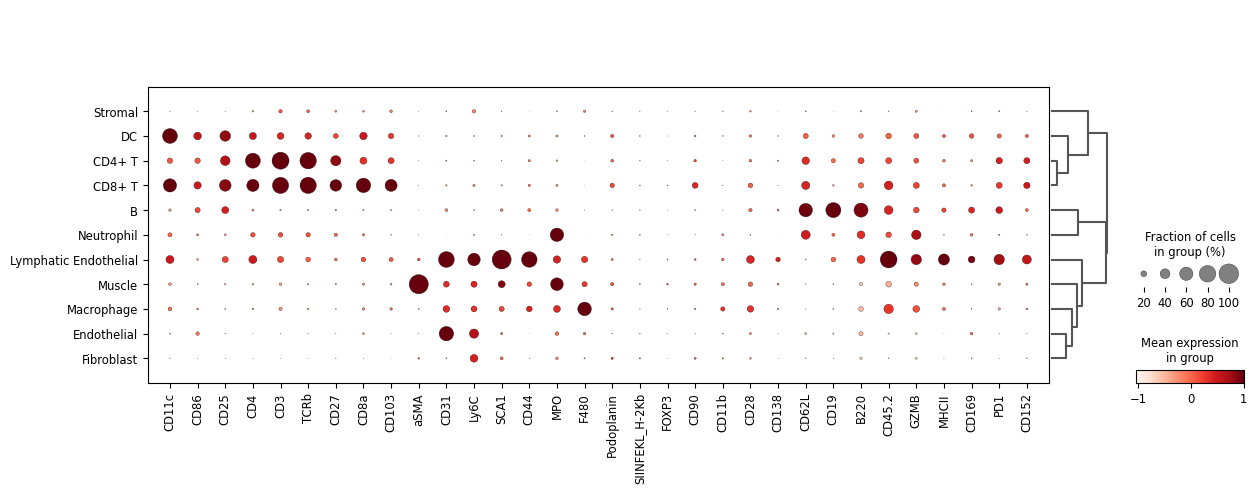

Supplementary Figure 5. Cell-type annotation in the multiplexed in vivo NanoSTAMP experiment. Dot plot showing protein-marker expression across annotated splenic cell populations 24 h after intravenous administration of the pooled LNP library. Dot size represents the percentage of cells within each population with z-normalized marker expression above 0.7, and color indicates mean z-normalized expression. The dendrogram shows hierarchical relationships among cell populations based on their marker-expression profiles. Data represent four mice, with four spleen sections per mouse processed independently as technical replicates.

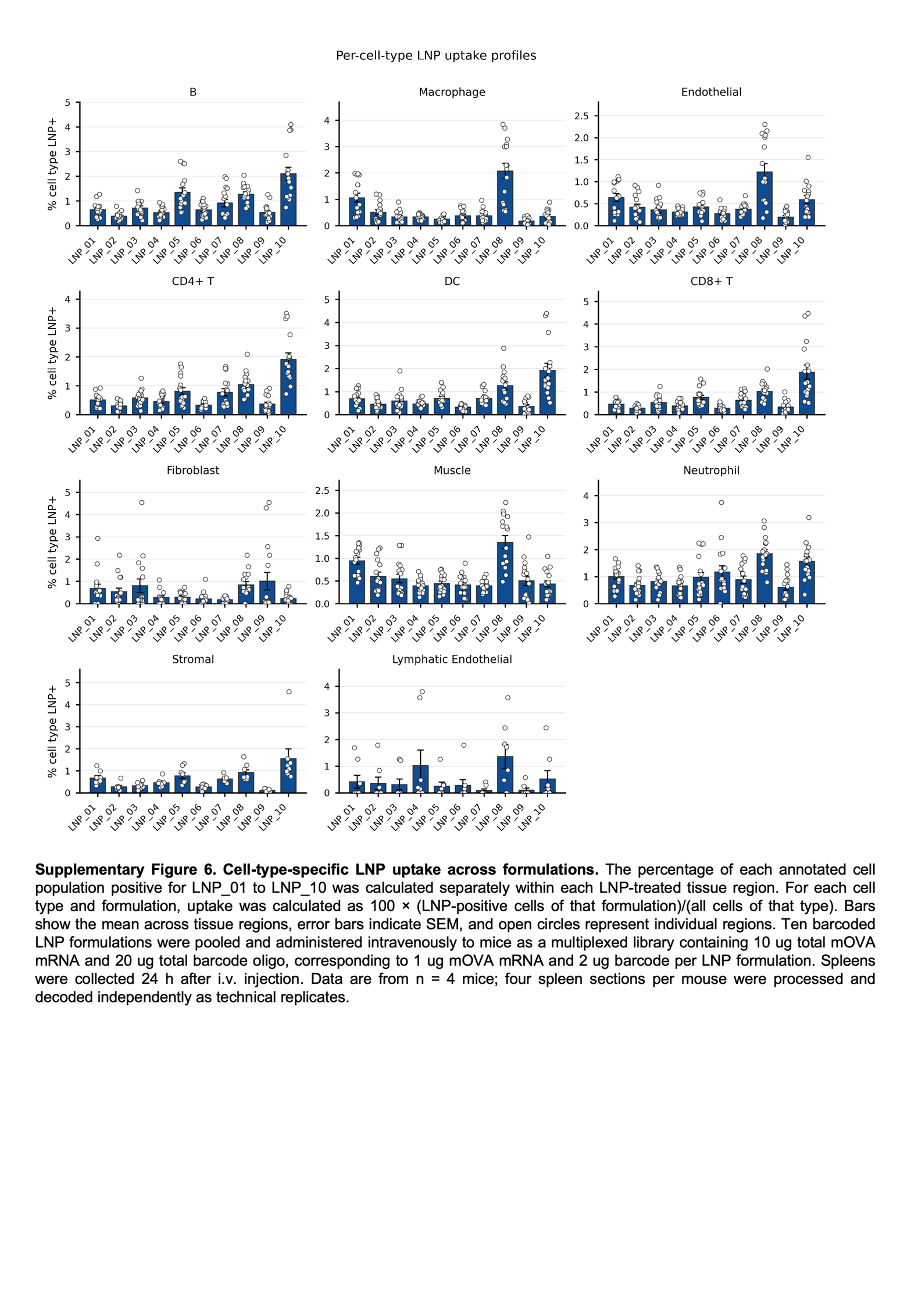

Supplementary Figure 6. Cell-type-specific LNP uptake across formulations. The percentage of each annotated cell population positive for LNP_01 to LNP_10 was calculated separately within each LNP-treated tissue region. For each cell type and formulation, uptake was calculated as 100 × (LNP-positive cells of that formulation)/(all cells of that type). Bars show the mean across tissue regions, error bars indicate SEM, and open circles represent individual regions. Ten barcoded LNP formulations were pooled and administered intravenously to mice as a multiplexed library containing 10 ug total mOVA mRNA and 20 ug total barcode oligo, corresponding to 1 ug mOVA mRNA and 2 ug barcode per LNP formulation. Spleens were collected 24 h after i.v. injection. Data are from n = 4 mice; four spleen sections per mouse were processed and decoded independently as technical replicates.

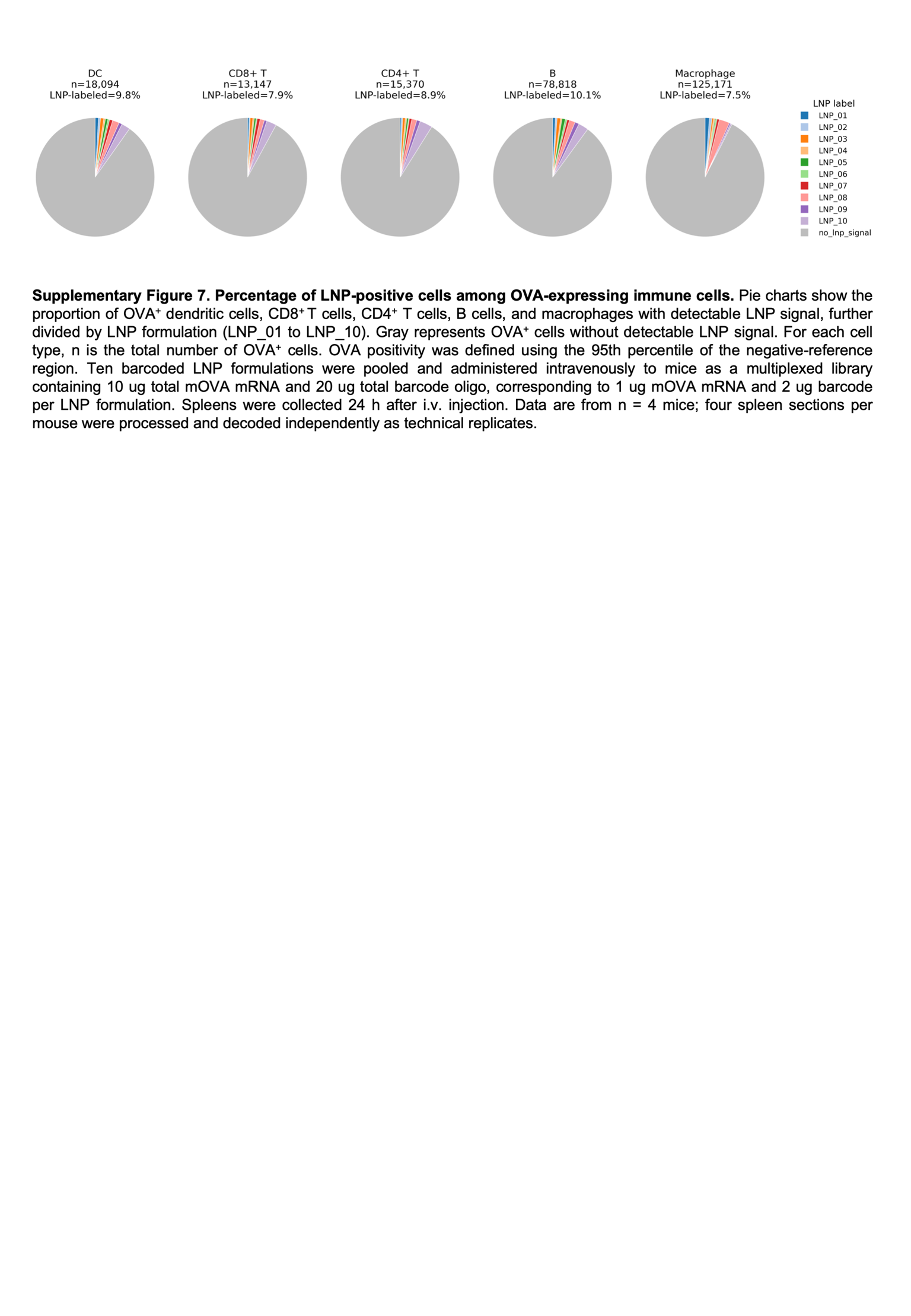

Supplementary Figure 7. Percentage of LNP-positive cells among OVA-expressing immune cells. Pie charts show the proportion of OVA^+^ dendritic cells, CD8^+^ T cells, CD4^+^ T cells, B cells, and macrophages with detectable LNP signal, further divided by LNP formulation (LNP_01 to LNP_10). Gray represents OVA^+^ cells without detectable LNP signal. For each cell type, n is the total number of OVA^+^ cells. OVA positivity was defined using the 95th percentile of the negative-reference region. Ten barcoded LNP formulations were pooled and administered intravenously to mice as a multiplexed library containing 10 ug total mOVA mRNA and 20 ug total barcode oligo, corresponding to 1 ug mOVA mRNA and 2 ug barcode per LNP formulation. Spleens were collected 24 h after i.v. injection. Data are from n = 4 mice; four spleen sections per mouse were processed and decoded independently as technical replicates.

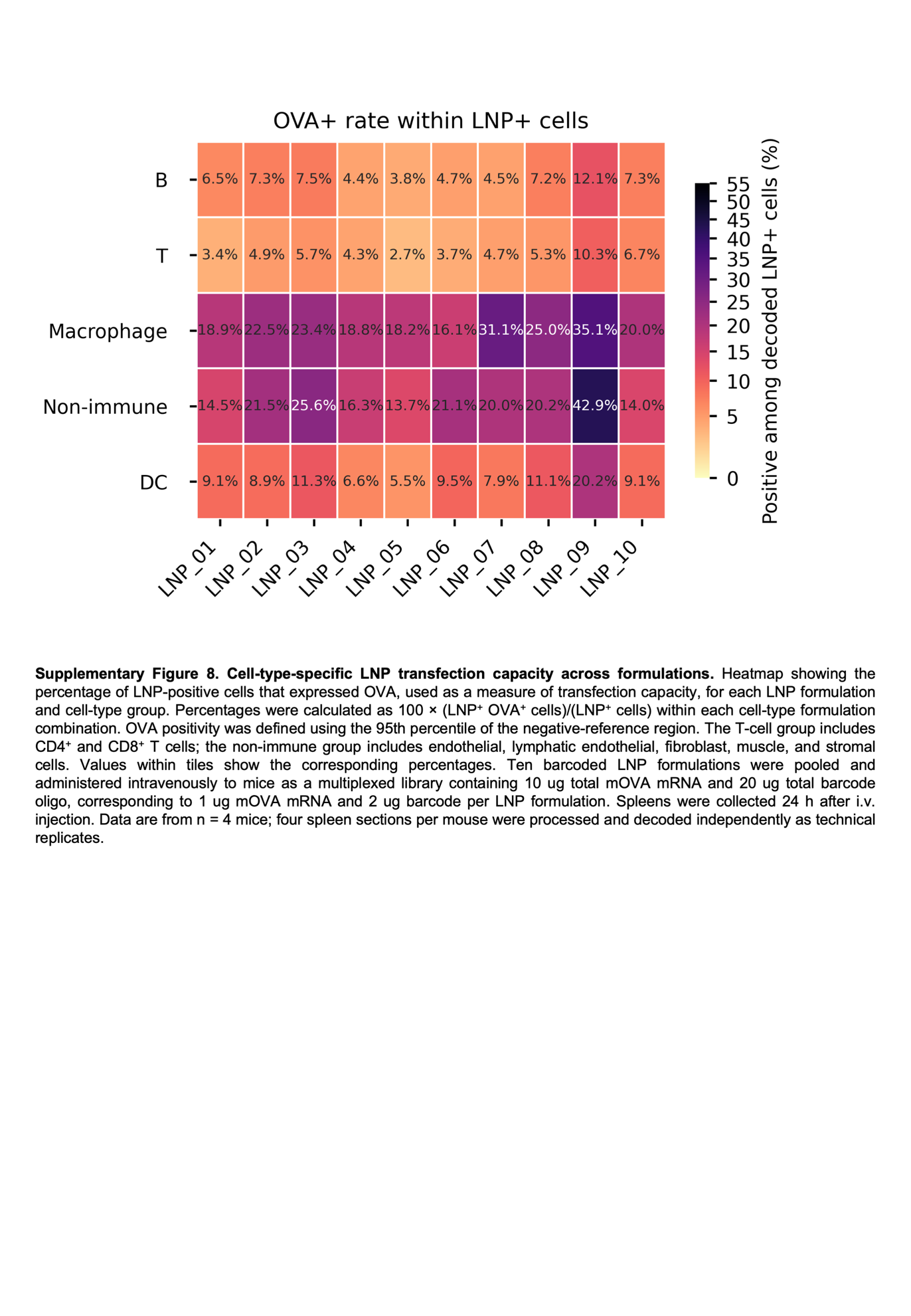

Supplementary Figure 8. Cell-type-specific LNP transfection capacity across formulations. Heatmap showing the percentage of LNP-positive cells that expressed OVA, used as a measure of transfection capacity, for each LNP formulation and cell-type group. Percentages were calculated as 100 × (LNP^+^ OVA^+^ cells)/(LNP^+^ cells) within each cell-type formulation combination. OVA positivity was defined using the 95th percentile of the negative-reference region. The T-cell group includes CD4^+^ and CD8^+^ T cells; the non-immune group includes endothelial, lymphatic endothelial, fibroblast, muscle, and stromal cells. Values within tiles show the corresponding percentages. Ten barcoded LNP formulations were pooled and administered intravenously to mice as a multiplexed library containing 10 ug total mOVA mRNA and 20 ug total barcode oligo, corresponding to 1 ug mOVA mRNA and 2 ug barcode per LNP formulation. Spleens were collected 24 h after i.v. injection. Data are from n = 4 mice; four spleen sections per mouse were processed and decoded independently as technical replicates.

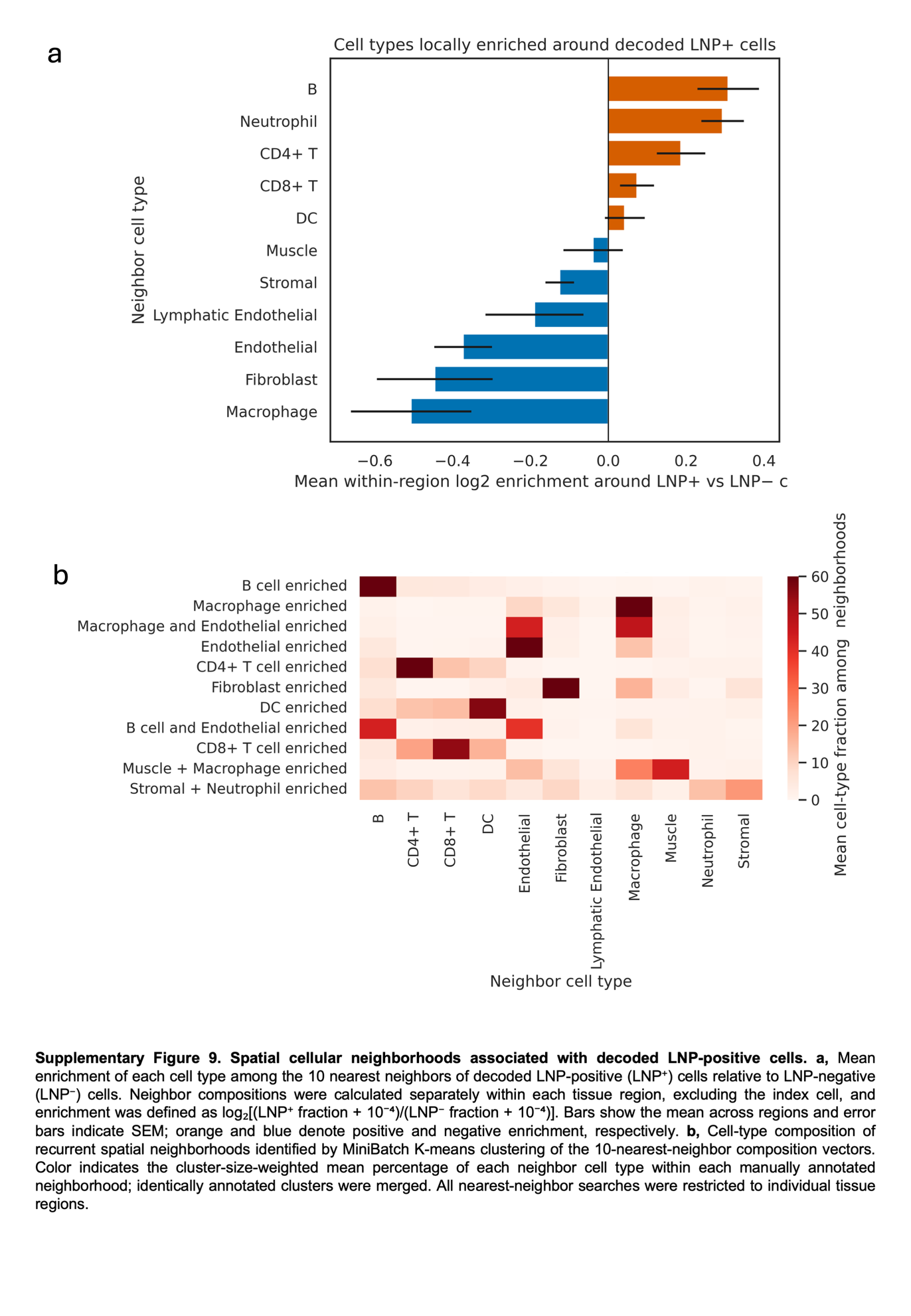

Supplementary Figure 9. Spatial cellular neighborhoods associated with decoded LNP-positive cells. (a), Mean enrichment of each cell type among the 10 nearest neighbors of decoded LNP-positive (LNP^+^) cells relative to LNP-negative (LNP^−^) cells. Neighbor compositions were calculated separately within each tissue region, excluding the index cell, and enrichment was defined as log₂[(LNP^+^ fraction + 10⁻⁴)/(LNP^−^ fraction + 10⁻⁴)]. Bars show the mean across regions and error bars indicate SEM; orange and blue denote positive and negative enrichment, respectively. (b), Cell-type composition of recurrent spatial neighborhoods identified by MiniBatch K-means clustering of the 10-nearest-neighbor composition vectors. Color indicates the cluster-size-weighted mean percentage of each neighbor cell type within each manually annotated neighborhood; identically annotated clusters were merged. All nearest-neighbor searches were restricted to individual tissue regions.

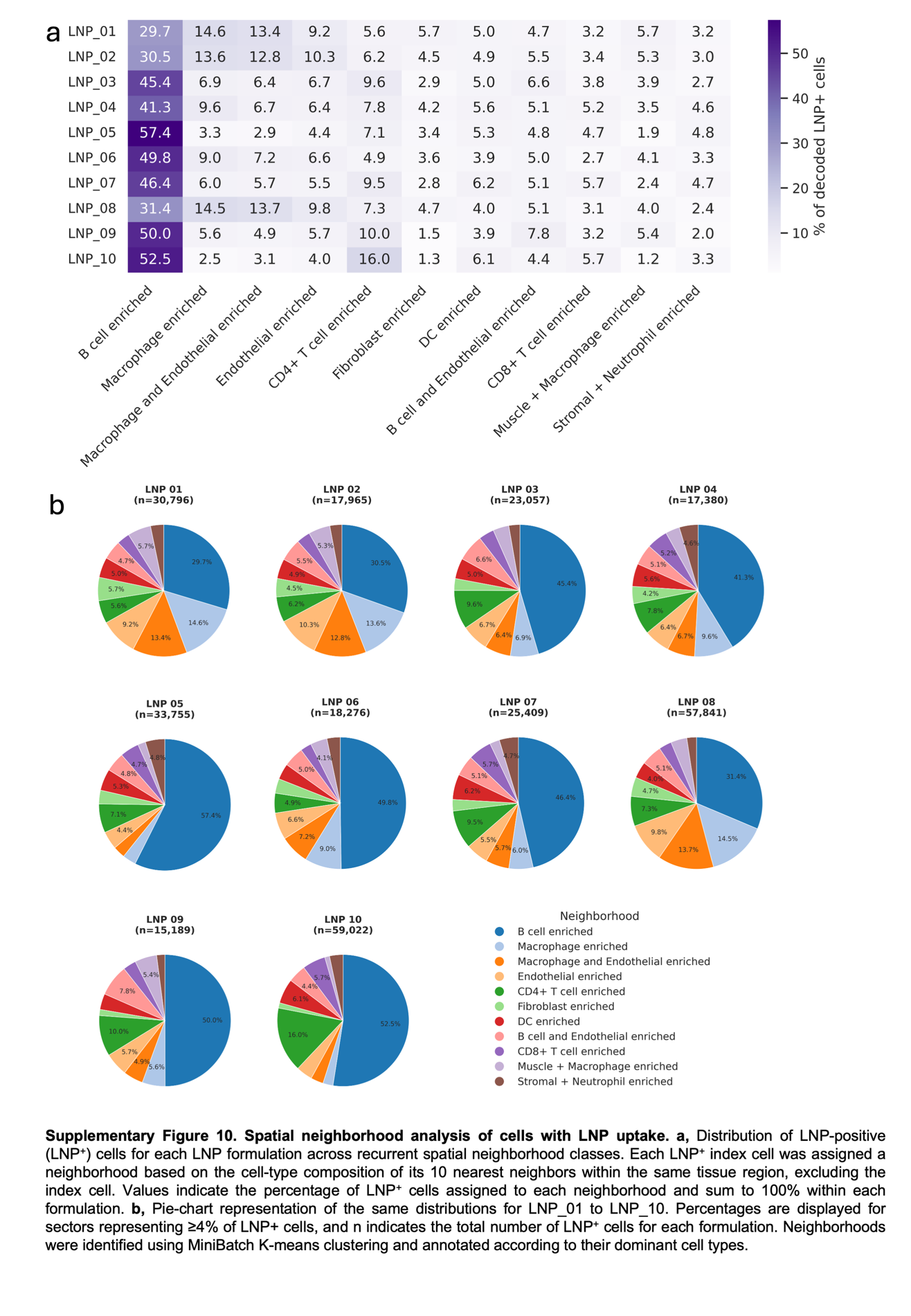

Supplementary Figure 10. Spatial neighborhood analysis of cells with LNP uptake. (a), Distribution of LNP-positive (LNP+) cells for each LNP formulation across recurrent spatial neighborhood classes. Each LNP+ index cell was assigned a neighborhood based on the cell-type composition of its 10 nearest neighbors within the same tissue region, excluding the index cell. Values indicate the percentage of LNP+ cells assigned to each neighborhood and sum to 100% within each formulation. (b), Pie-chart representation of the same distributions for LNP_01 to LNP_10. Percentages are displayed for sectors representing ≥4% of LNP+ cells, and n indicates the total number of LNP+ cells for each formulation. Neighborhoods were identified using MiniBatch K-means clustering and annotated according to their dominant cell types.

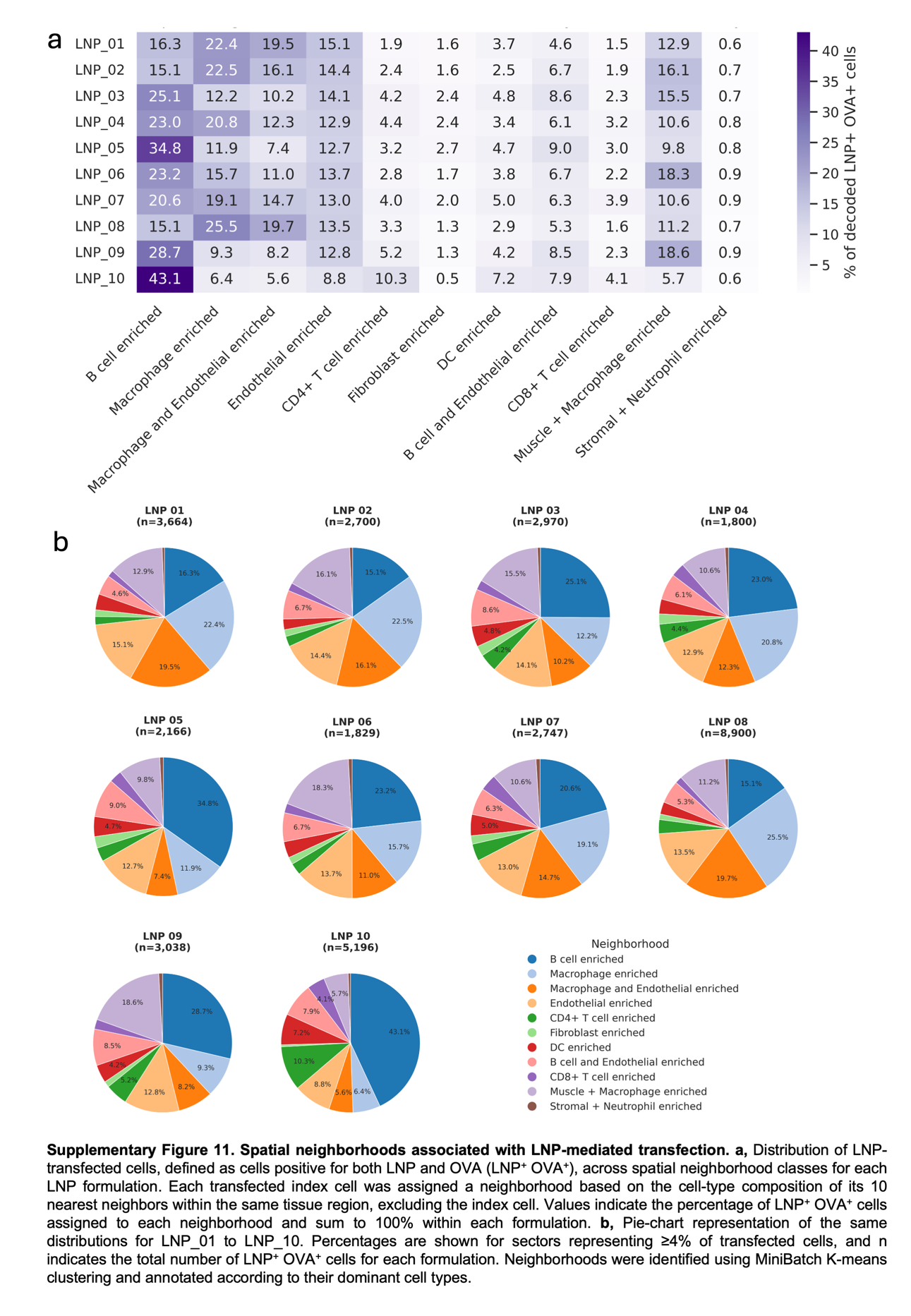

Supplementary Figure 11. Spatial neighborhoods associated with LNP-mediated transfection. (a), Distribution of LNP-transfected cells, defined as cells positive for both LNP and OVA (LNP^+^ OVA^+^), across spatial neighborhood classes for each LNP formulation. Each transfected index cell was assigned a neighborhood based on the cell-type composition of its 10 nearest neighbors within the same tissue region, excluding the index cell. Values indicate the percentage of LNP^+^ OVA^+^ cells assigned to each neighborhood and sum to 100% within each formulation. (b), Pie-chart representation of the same distributions for LNP_01 to LNP_10. Percentages are shown for sectors representing ≥4% of transfected cells, and n indicates the total number of LNP^+^ OVA^+^ cells for each formulation. Neighborhoods were identified using MiniBatch K-means clustering and annotated according to their dominant cell types.

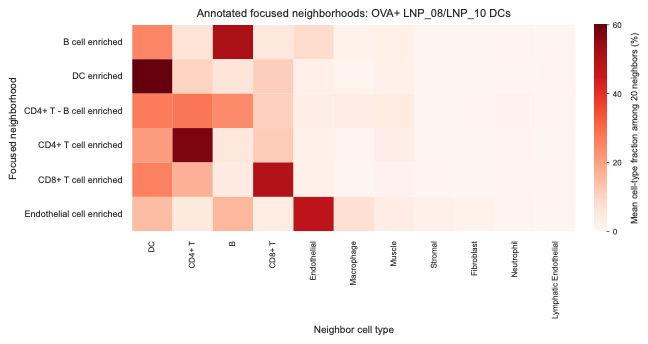

Supplementary Figure 12. Focused spatial neighborhoods of OVA⁺ LNP⁺ dendritic cells. Heatmap showing the mean cell-type composition of the 20 nearest non-self neighbors (k=20) for neighborhood classes identified among pooled LNP08- and LNP10-associated OVA⁺ dendritic cells. Rows indicate annotated neighborhood classes, columns indicate neighboring cell types.

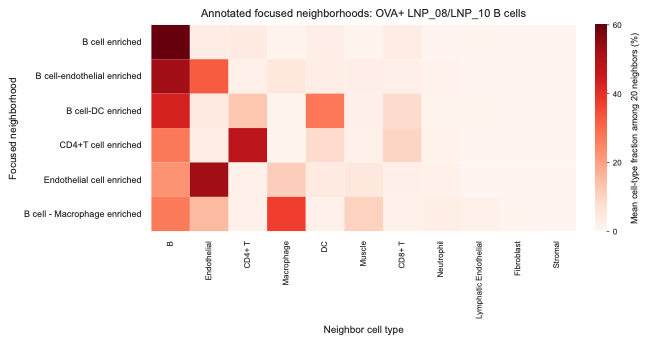

Supplementary Figure 13. Focused spatial neighborhoods of OVA⁺ LNP⁺ B cells. Heatmap showing the mean cell-type composition of the 20 nearest non-self neighbors (k=20) for neighborhood classes identified among pooled LNP08- and LNP10-associated OVA⁺ B cells. Rows indicate annotated neighborhood classes, columns indicate neighboring cell types.

**Supplementary Table 1. Delivery oligonucleotide and padlock probe sequences used in NanoSTAMP.**

| **Oligo set** | **Delivery oligonucleotide sequence (5′–3′)** | **Padlock probe sequence (5′–3′)** |
| --- | --- | --- |
| **Single_test** | GCCGTAGTGT TCATGAATGT AGGATGGTCT CCACTGATTA AGAGGCG | /5Phos/ TCATGAACAC TACGGCATCG TAACACATCC ATAACGACCA TGTGACCTTG ACGACCGACA GTGAGACCAT CCTACATTGT AGTCCTAAAC CATTACTCTC TTAAGCTAAT TCGCCTCTTA ATCAGTG |
| 10plex_pool_1 | TAGGAGTGTA ATAGCCGGCG TCATTCCCGG AGTCTACCCA ACGCAGGCAG TCTACTACCT | /5Phos/ CCGGGAATGA CGCCGGCTAT TACACTCCTA TAATCGTAAC ACATCCATGT GCGAGTTGAC CTTACCACTT CCGGCTGAGA CCATCCTACA TTGTAGTCCT AAACCATTAC TCTCTTAAGC TAATTTAAGG TAGTAGACTG CCTGCGTTGG GTAGACT |
| 10plex_pool_2 | CACCAAGTCT GTAGAACTCC GTTCTTCTCT TCTACGTGCG AGCTCAACAC CATGATTGCG | /5Phos/ AGAAGAGAAG AACGGAGTTC TACAGACTTG GTGTAATCGT AACACATCCA TAACGACCAT GTGACCTTGA CGACCGACAG TGAGACCATC CTACATTGTA GTCCTAAACC ATTACTCTCT TAAGCTAATT TACGCAATCA TGGTGTTGAG CTCGCACGT |
| 10plex_pool_3 | TGCTCGTAGC ACGCAGGGAC TTTCGCTAAT TATGAACTGG TACGGTTCAC TCCCAACCTA | /5Phos/ AATTAGCGAA AGTCCCTGCG TGCTACGAGC ATAATCGTAA CACATCCATG TGCGAGTTGA CCTTTGACGA CCGACAGTTA CGTTAGTGGA CCATTGTAGT CCTAAACCAT TACTCTCTTA AGCTAATTTA TAGGTTGGGA GTGAACCGTA CCAGTTCAT |
| 10plex_pool_4 | AGGGTACGAA GGGTGTAGAG ATTGGAGAGT ATAGCTACAT CGCGAAGAAC CGGCCTGTGT | /5Phos/ TCTCCAATCT CTACACCCTT CGTACCCTTA AACGACCATG TGACCTACCA CTTCCGGCTG AGACCATCCT ACATTTACGT TAGTGGACCA TATCCGTCGC GTTTGTAGTC CTAAACCATT AACACAGGCC GGTTCTTCGC GATGTAGCTA TAC |
| 10plex_pool_5 | TAGCTTTGTC CGGCGCTTCG CTTCTTTACG CAACCATTGC GCATCATAGG GTGCATGTTA | /5Phos/ CGTAAAGAAG CGAAGCGCCG GACAAAGCTA TAAAGCGGTC TTACGGTTGT GCGAGTTGAC CTTTGACGAC CGACAGTGAG ACCATCCTAC ATTATCCGTC GCGTTTACTC TCTTAAGCTA ATTTATAACA TGCACCCTAT GATGCGCAAT GGTTG |
| 10plex_pool_6 | TTTCCAATCG TCGTCATTTA CCGGTGACCG ATCATCTGCG TAATCTGGTA TCGTCTGCAC | /5Phos/ TCGGTCACCG GTAAATGACG ACGATTGGAA ATAAAGCGGT CTTACGGTTG AGACCATCCT ACATTCTCAG GTTCGAGTCT ATCCGTCGCG TTTGTAGTCC TAAACCATTA CTCTCTTAAG CTAATTTAGT GCAGACGATA CCAGATTACG CAGATGA |
| 10plex_pool_7 | ATAGCCTCAA GTGACGAAGT CGTGCTAGCT CTACAAATAC CCGGATTTGA CGAGAGTCCT | /5Phos/ GAGCTAGCAC GACTTCGTCA CTTGAGGCTA TTAAAGCGGT CTTACGGTTG TGCGAGTTGA CCTTAACGAC CATGTGACCT TGACGACCGA CAGTCTCAGG TTCGAGTCTA TCCGTCGCGT TTAAGGACTC TCGTCAAATC CGGGTATTTG TA |
| 10plex_pool_8 | CTTGTGTGCT TAGACCTGGG AATGTCGAGG CCTCGCGTCG AACTCATTGT GCTATGCCGA | /5Phos/ CCTCGACATT CCCAGGTCTA AGCACACAAG TAATCGTAAC ACATCCATAA GCGGTCTTAC GGTTGTGCGA GTTGACCTTA ACGACCATGT GACCTACCAC TTCCGGCTCT CAGGTTCGAG TCTATCGGCA TAGCACAATG AGTTCGACGC GAGG |
| 10plex_pool_9 | TGATTTCTCG TTGAACCACG AGGTAATCGA TGGATAGACA GTATGGTCAC GCCCAGTGAA | /5Phos/ CGATTACCTC GTGGTTCAAC GAGAAATCAT AAAGCGGTCT TACGGTTAAC GACCATGTGA CCTCTCAGGT TCGAGTCTTA CGTTAGTGGA CCATTGTAGT CCTAAACCAT TACTCTCTTA AGCTAATTTA TTCACTGGGC GTGACCATAC TGTCTATCCA T |
| 10plex_pool_10 | TCGTTACCAG CGACACTCGC TGGCTTGATG CCCGTCATAG ACGAATCGAA GACCTCCGAA | /5Phos/ CATCAAGCCA GCGAGTGTCG CTGGTAACGA TAAAGCGGTC TTACGGTTGT GCGAGTTGAC CTTTGACGAC CGACAGTACC ACTTCCGGCT ATCCGTCGCG TTTGTAGTCC TAAACCATTA TTCGGAGGTC TTCGATTCGT CTATGACGGG |

* The Single_test oligonucleotide was used for all in vitro experiments and the single-formulation SM-102 in vivo experiment. The 10plex_pool_1–10 delivery oligonucleotides and corresponding padlocks were used for the multiplexed in vivo study of ten LNP formulations. All sequences are shown 5′–3′. /5Phos/ indicates a 5′ phosphate modification. Spaces within sequences are included only for readability.

**Supplementary Table 2. CODEX protein panel staining conditions and cycle information**

| Cycles | Ax488 | Oligo | µL/ slide | Exp. time (ms) | Cy3 | Oligo | µL/ slide | Exp. time (ms) | Cy5 | Oligo | µL/ slide | Exp. time (ms) |
| --- | --- | --- | --- | --- | --- | --- | --- | --- | --- | --- | --- | --- |
| 1 | aSMA | 69 | 1 | 150 | Podoplanin | 74 | 2 | 150 | CD4 | 7 | 1 | 150 |
| 2 | SCA1 | 14 | 2 | 150 | CD31 | 60 | 0.5 | 150 | NKp46 | 45 | 1 | 150 |
| 3 | Ly6C | 41 | 0.5 | 150 | CD45.2 | 51 | 2 | 150 | PD1 | 23 | 1 | 150 |
| 4 | Blank |  |  |  | CD3 | 81 | 1 | 150 | CD11c | 26 | 1 | 150 |
| 5 | Blank |  |  |  | CD19 | 24 | 1 | 150 | F480 | 77 | 2 | 150 |
| 6 | Blank |  |  |  | CD138 | 53 | 4 | 150 | CD169 | 59 | 2 | 150 |
| 7 | Blank |  |  |  | CD62L | 43 | 1 | 150 | TCRb | 3 | 0.4 | 150 |
| 8 | Blank |  |  |  | GZMB | 57 | 1 | 150 | FOXP3 | 61 | 2 | 150 |
| 9 | Blank |  |  |  | CD27 | 32 | 1 | 150 | CD8a | 8 | 1 | 150 |
| 10 | Blank |  |  |  | CD11b | 5 | 0.5 | 150 | FLuc | 11 | 4 | 150 |
| 11 | Blank |  |  |  | MHCII | 36 | 0.5 | 150 | CCR7 | 62 | 1 | 150 |
| 12 | Blank |  |  |  | CD86 | 66 | 1 | 150 | CD28 | 48 | 4 | 150 |
| 13 | Blank |  |  |  | B220 | 71 | 2 | 150 | CD103 | 29 | 1 | 150 |
| 14 | Blank |  |  |  | CD25 | 68 | 4 | 150 | MPO | 58 | 2 | 150 |
| 15 | Blank |  |  |  | CD90 | 70 | 2 | 150 | CD152 | 42 | 1 | 150 |
| 16 | Blank |  |  |  | CD44 | 44 | 0.5 | 150 | SIINFEKL H-2Kb | 21 | 4 | 150 |
| 17 | Blank |  |  |  |  |  |  |  | OVA | 15 | 4 | 150 |
| 0 | Reference |  |  | 6 | Reference |  |  | 6 | Reference |  |  | 6 |

**Supplementary Table 3. CODEX post-RCA staining conditions and cycle information**

| Cycles | Ax488 | Oligo | µL/ slide | Exp. time (ms) | Cy3 | Oligo | µL/ slide | Exp. time (ms) | Cy5 | Oligo | µL/ slide | Exp. time (ms) |
| --- | --- | --- | --- | --- | --- | --- | --- | --- | --- | --- | --- | --- |
| 1 | Blank |  |  |  | Blank |  |  |  | Bit_1 | 6 | NA | 200 |
| 2 | Blank |  |  |  | Blank |  |  |  | Bit_2 | 17 | NA | 200 |
| 3 | Blank |  |  |  | Blank |  |  |  | Bit_3 | 55 | NA | 200 |
| 4 | Blank |  |  |  | Blank |  |  |  | Bit_4 | 56 | NA | 200 |
| 5 | Blank |  |  |  | Blank |  |  |  | Bit_5 | 76 | NA | 200 |
| 6 | Blank |  |  |  | Blank |  |  |  | Bit_6 | 79 | NA | 200 |
| 7 | Blank |  |  |  | Blank |  |  |  | Bit_7 | 20 | NA | 200 |
| 8 | Blank |  |  |  | Blank |  |  |  | Bit_8 | 46 | NA | 200 |
| 9 | Blank |  |  |  | Blank |  |  |  | Bit_9 | 28 | NA | 200 |
| 10 | Blank |  |  |  | Blank |  |  |  | Bit_10 | 72 | NA | 200 |
| 11 | Blank |  |  |  | Blank |  |  |  | Bit_11 | 2 | NA | 200 |
| 12 | Blank |  |  |  | Blank |  |  |  | Bit_12 | 63 | NA | 200 |
| 0 | Reference |  |  | 6 | Reference |  |  | 6 | Reference |  |  | 6 |

**Supplementary Table 4. CODEX antibody information**

| RRID | Company | Antibody | Clone | RRID | Company | Antibody | Clone |
| --- | --- | --- | --- | --- | --- | --- | --- |
| AB_2572996 | Invitrogen | aSMA | 1A4 | AB_394606 | BD | CD45 | 30-F11 |
| AB_1107651 | BioXcell | B220 | Ra3-6B2 | NA | SinoBiological | CD62L | 414 |
| AB_389229 | BioLegend | CCR7 | 4B12 | AB_313144 | BioLegend | CD86 | GL-1 |
| AB_535944 | BioLegend | CD103 | 2E7 | AB_2275792 | BD | CD8a | 53-6.7 |
| AB_393577 | BD | CD11b | M1/70 | AB_313168 | BioLegend | CD90 | G7 |
| AB_313770 | BioLegend | CD11c | N418 | AB_2869866 | BD | F4/80 | T45-2342 |
| AB_313251 | BioLegend | CD152 | UCT10-4B9 | AB_467576 | eBioscience | FOXP3 | FJK-16s |
| NA | BioRad | CD169 | MOMA-1 | NA | Novus | GRZB | AF1865 |
| AB_395047 | BD | CD19 | 1D3 | AB_1877086 | BioLegend | PD1 | 29F.1A12 |
| AB_1236456 | BioLegend | CD27 | LG.3A10 | AB_1089187 | BioXcell | Podoplanin | 8.1.1 |
| AB_1107624 | BioXcell | CD28 | 37.51 | AB_313348 | BioLegend | Sca-1 | D7 |
| AB_395697 | BD | CD3 | 17A2 | AB_10950158 | BioXcell | TCRb | H57-597 (HB218) |
| AB_394815 | BD | CD31 | MEC 13.3 | AB_2539921 | Thermo | OVA | poly |
| AB_393575 | BD | CD4 | RM4-5 | AB_925771 | invitrogen | OVA257-264 | eBio25-D1.16 |
| AB_394645 | BD | CD44 | IM7 | AB_2549682 | Thermo | Luciferase | poly |
| AB_1134214 | BioLegend | Ly6C | HK1.4 | NA | BioXcell | MPO | 6G4 |
| AB_313316 | BioLegend | MHCII | M5/114.15.2 | AB_1727467 | BD | NKp46 | 29A1.4 |
| AB_10959655 | Biolegend | CD138 | 281-2 | AB_312851 | Biolegend | CD25 | PC61 |
